# Higher Order AMMI (HO-AMMI) analysis: A novel stability model to study genotype-location interactions

**DOI:** 10.1101/2020.10.05.326538

**Authors:** BC Ajay, Fiyaz R Abdul, SK Bera, Narendra kumar, K Gangadhar, Praveen kona, Kirti Rani, T Radhakrishnan

## Abstract

Additive main effects and multiplicative interaction (AMMI) model is most widely used to analyse genotype*environment interactions (GEI) wherein interaction effects of location is masked by year effect. Hence, presently available models are not able to estimate interaction effects of genotype*location (GLI) and genotype*year (GYI) separately. Moreover, genotype ranking differs as number of years of evaluation vary making selection of genotype for target location difficult. In the present study we propose a novel stability model i.e Higher-order-AMMI (HO-AMMI) analysis which is capable of calculating GLI without the confounding effect of GYI and GLYI. GEI of AMMI model and all 2-way interactions of HO-AMMI model follow χ^2^ distribution, whereas 3-way interaction (GLYI) of HO-AMMI follow noncentral χ^2^ distribution. With increase in number of years of evaluation contribution of GLI towards total variation increased whereas in AMMI model contribution of GEI towards total variation decreased. Variation explained by multiplicative components is higher in HO-AMMI compared to AMMI model. Genotypes were ranked using GL, GY and GL+GY+GLY interactions of HO-AMMI and GEI of AMMI for stability and yield and compared their ranks with field ranking. Correlation and linear regression analysis have indicated high association of GLI (HO-AMMI) with field ranking with high R^2^ values. Further, HO-AMMI model was able to remove the confounding effect of GYI and GLYI on GLI for accurate identification of genotype for target location irrespective of number of years of evaluation. Hence, HO-AMMI model can be used under multi-environment trials (MET) for selecting genotypes efficiently.

**Availability and Information:** Source code implemented in R is available from corresponding author.

## 1. Introduction

Often it is assumed that the phenotype (Y) is the sum of the genotype (G) and the environment (E) (Y=G+E). The phenotype can also be seen as the interaction between genotypical and environmental factors, and some genotypes are better adapted to one environment and others are better in another environment. If a genotype performs well in one environment, the same genotype will not necessarily result in a good phenotype in another environment. Therefore, phenotype was described as a function of the genotype and the environment. As per the Fisherian quantitative genetics, observed phenotype (Y) is due to the combined action of genotype (G), environmental (E) effects plus genotype*environment interaction (GE) effects:

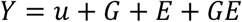

Where ‘u’ is population mean. Means across environments are adequate indicators of genotypic performance of trials when GE interactions are non-significant. But, when GE interactions are significant, means mask subsets of environments where genotype differ significantly in relative performance (Fox et al. 1977). Genotype selected in one environment/location may perform poorly in another environment/location. As a result, plant breeders often consider performance of genotype across environments/locations. Hence, statistical analysis of multi-environment trials (METs) should detect GE interactions, quantify it so as to identify adaptable genotype. Most of the MET trials involve evaluating genotypes under different locations (L) and years (Y). Then GE term in analysis of variance can be partitioned into location*year (LY) interaction, genotype*location (GL) interactions, genotype*year (GY) interaction and genotype*location*years (GLY) interaction. Then, the above phenotypic expression is modified into:

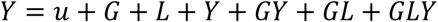

When GL is the dominant portion of GE, then specific adaptation can be exploited by subdividing the locations into homogenous regions that minimise GE within the regions. But when GY and GLY terms dominate, subdivision of locations is not possible to minimise GE and in such a cases representative location should be considered to determine genotypic responses (Fox et al. 1997). GE interaction effects play an important role in phenotypic expression of a trait and can be studied using several methods, including linear-bilinear models. The additive main effects and multiplicative interaction (AMMI) model is currently one of the most popular multiplicative models. Though AMMI model was initially proposed by Gollob (1968) for fixed effects, but the actual statistical methodology was credited to the works of Pike and Silverberg (1952) and Williams (1952). It has been widely used to study GE interactions in plant breeding programs and in agronomic trials. AMMI model is fit in two stages. First, main effects of the model are estimated using the additive two-way analysis of variance (ANOVA) by least squares. Then, the singular value decomposition (SVD) is applied to the residuals from the ANOVA, i.e. to the interaction, to obtain the estimates for the multiplicative terms of the AMMI model (Gauch 1988). AMMI analysis has been used to study the stability of genotypes over environments in several crops (Ajay et al. 2019; Chuni Lal et al. 2019; Bhartiya et al. 2017). Caliskan et al. (2007) used AMMI model to study GE interactions in eleven sweetpotato genotypes tested over eight environments i.e four locations during 2 years. Similarly, Madry et al. (2017) studied the response of winter wheat cultivars to crop management and environments using AMMI model. Here, 24 wheat cultivars were tested at two crop management intensities, 20 locations and three years (2006-2008) and yield data were analysed by two stage approach. In the first, stage, regular ANOVA was performed considering all four factors (i.e genotypes, crop management, location and year) and studied all possible two-way and three-way interactions. In the second stage, while performing AMMI model, they computed interaction principal components (IPCAs) only for genotype* environment interactions (GEI) considering each location and year as one environment. As described by Fox et al. (1997) GEI may be further partitioned into genotype*location interaction (GLI), genotype*year interaction (GYI) and genotype*location*years interaction (GLYI) when location and year factor is involved. Currently available AMMI models compute only GEI considering location and year as one environment and none of the currently available models partition the GEI into GLI, GYI and GLYI.

Hence, the objective of the present study was to propose a novel stability model referred as “Higher-Order Additive Main effects and Multiplicative Interaction (HO-AMMI)” model which can decompose GEI into GYI, GLI and GLYI and compute IPCAs for GLI alone without the confounding effects of GYI and GLYI. Further we compute AMMI stability value using Modified AMMI stability Index (MASI) (Ajay et al. 2018a) by using IPCAs from GYI, GLI and GLYI. We further compare the results of HO-AMMI and AMMI models to show their efficiency in ranking genotypes.

## 2. Material and methods

### 2.1 Higher Order-Additive Main effects and Multiplicative Interaction (HO-AMMI)

HO-AMMI model computes all possible two-way and three-way interactions such as GLI, GYI, GLYI separately to calculate stability values without the confounding effects of other interactions. IPCAs estimated using GLI could be used to identify genotypes for target location precisely. Higher-Order AMMI model equation can be written as;

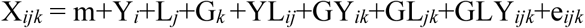

Where,

X_ijk_ – Yield in i^th^ year, j^th^ location and k^th^ genotype

m – General mean

Y_i_ – random main effect of i^th^ year

L_j_ – fixed main effect of j^th^ location

G_k_ – fixed main effect of k^th^ genotype

YL_ij_ – random interaction effect of i^th^ year and j^th^ location

GY_ik_ – random interaction effect of k^th^ genotype and i^th^ year

GL_ik_ – random interaction effect of k^th^ genotype and j^th^ location

GLY_ijk_ – random interaction effect of k^th^ genotype, in i^th^ year and j^th^ location

e_ijk_ – average error associated with the response of the k^th^ genotype in i^th^ year and j^th^ location

In HO-AMMI model, all two-way (GYI, GLI and LYI) and three-way (GLYI) interactions were computed separately following “factor analytic model” suggested by Gollob (1968) and Gauch (1988) using SVD. HO-AMMI model involving SVD can be represented as,

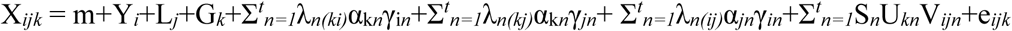

Where,

λ_*n(ki)*_ – singular value of n^th^ multiplicative component for k^th^ genotype in i^th^ year

λ_*n(kj)*_ – singular value of n^th^ multiplicative component for k^th^ genotype in j^th^ location

λ_*n(ij)*_ – singular value of n^th^ multiplicative component for i^th^ year in j^th^ location

S_n_ – singular value of n^th^ multiplicative component for k^th^ genotype in i^th^ year and j^th^ location

α_k*n*_ – n^th^ singular vector for k^th^ genotypes

α_j*n*_ – n^th^ singular vector for j^th^ location

γ_i*n*_ –n^th^ singular vector for i^th^ year

γ_*jn*_ – n^th^ singular vector for j^th^ location

U_*kn*_ – combined n^th^ singular vector for k^th^ genotypes

V_*ijn*_ – combined n^th^ singular vector for i^th^ year in j^th^ location

### 2.2 Simulated data and comparison of models

In order to compare AMMI and HO-AMMI models, simulated data of peanut pod yield were generated for four different experiments. All four experiments consisted of 20 locations uniformly. Experiments 1 and 2 had 10 genotypes with 2 and 3 years of evaluation whereas experiments 3 and 4 had 20 genotypes each with 2 and 3 years of evaluation respectively. AMMI model analysis was performed separately for four experiments in R (R core team, 2018) using package ‘agricolae’ (de Mendiburu 2017) and further Modified AMMI stability Index (MASI) was calculated as described by Ajay et al. (2018a) using the package ‘ammistability’ (Ajay et al. 2018b). HO-AMMI analysis was performed in R (R core team 2018) using packages ‘MASS’ (Ripley et al. 2019) and further MASI was calculated using SVDs from GY, GL and GLY interactions using the package ‘ammistability’ (Ajay et al. 2018b). Then, simultaneous selection index for yield and stability (SSI) was calculated using genotype ranking based on MASI and pod yield for both AMMI and HO-AMMI models. SSI rankings were correlated with field ranking using spearman’s rank correlation (Spearman, 1904) for model comparison.

### 2.3 Field Ranking

In order to arrive at field ranking of genotypes, every location and year in an experiment were considered as separate environments. For example, in experiment-1, 20 location and 2 years were considered as 40 environments and genotypes were ranked for all environments separately. Number of times genotype received rank ‘1’ over 40 environments was computed. Similarly, number of times genotype receiving ranks ‘2’ to ‘10’ were computed. In case of experiments 3 and 4 with 20 genotypes number of times a genotype receiving ranks ‘1’ to ‘20’ were computed. Overall genotype ranking was worked out using Garrett’s ranking method (Garratt & Woodworth, 1971) with the help of following formula,

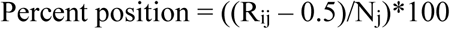

Where, R_ij_ = Rank given for i^th^ genotype for j^th^ rank, N_j_ = number of genotypes ranked.

Percent position for each genotype is converted into scores by using Garrett’s Table. Then, for each genotype, scores of all different ranks were added to get total score value and mean value of scores was calculated. Genotype having highest mean score value gets first rank and genotype with least mean score value gets last rank.

### 2.4 Model comparison

In order to check the efficiency of HO-AMMI model, spearman’s rank correlation was performed between GLI, GYI, GLYI and GLI+GYI+GLYI from HO-AMMI and GEI from AMMI model with field ranking in PAST statistical software (Hammer et al. 2001) to identify the model best associated with field ranking. Later, linear regression plots with R^2^ values were drawn for GLI of HO-AMMI and GEI of AMMI models to identify the model with high R^2^ values. Root Mean Square error (RMSE) values were calculated to compare the efficiency between GLI of HO-AMMI and GEI of AMMI models.

## 3 Results and Discussion

### 3.1 Additive Main effect and Multiplicative Interaction (AMMI)

Table 1 presents the AMMI analysis of variance for simulated peanut pod yield data set from four different experiments. In this model sources of variation were grouped into main effects such as genotype (G) and environment (E) and interaction effect such as genotype*environment (GEI). All three sources of variation significantly influenced pod yield among four experimental data sets. Environment had maximum variation i.e 73.19%, 75.74%, 70.57% and 72.87% respectively in experiment-1, 2, 3 and 4 followed by GEI explaining 22.41%, 20.37%, 22.21% and 20.20% variation respectively in experiment-1, 2, 3 and 4 whereas G had least influence among all four experiments. In experiment 1 and 2, nine IPCAs were significant explaining 100% variation in GEI sum of squares whereas ten IPCAs were significant in experiment 3 and 4 explaining 100% variation in GE sum of squares in their respective experiments.

**Table 1:** AMMI analysis of variance for studying genotype-environment-interactions (GEI) under different combinations of genotype, location and year

|  | Experiment-1 |  |  | Experiment-2 |  |  | Experiment-3 |  |  | Experiment-4 |  |  |
| --- | --- | --- | --- | --- | --- | --- | --- | --- | --- | --- | --- | --- |
|  | Df | MSS | GEI<br>SS% | Df | MSS | GEI<br>SS% | Df | MSS | GEI<br>SS% | Df | MSS | GEI<br>SS% |
| Environment (E) | 39 | 23532482** | 73.19 | 59 | 25529364** | 75.74 | 39 | 40270269** | 70.57 | 59 | 43687365** | 72.87 |
| Genotype (G) | 9 | 803185** | 0.58 | 9 | 1370809** | 0.62 | 19 | 4096885** | 3.50 | 19 | 6992618** | 3.76 |
| Rep (E) | 80 | 52401 | 0.33 | 120 | 45568 | 0.27 | 80 | 89656 | 0.32 | 120 | 77967 | 0.26 |
| G*E | 351 | 800638** | 22.41 | 531 | 762875** | 20.37 | 741 | 667166** | 22.21 | 1121 | 637536** | 20.20 |
| IPCA1 | 47 | 2140297** | 35.80 | 67 | 2226431** | 36.82 | 57 | 3052948** | 35.2 | 77 | 3355380** | 36.15 |
| IPCA2 | 45 | 1160549** | 18.58 | 65 | 1154583** | 18.53 | 55 | 1637548** | 18.22 | 75 | 1725544** | 18.11 |
| IPCA3 | 43 | 832427** | 12.74 | 63 | 869859.3** | 13.53 | 53 | 1190451** | 12.76 | 73 | 1326417** | 13.55 |
| IPCA4 | 41 | 713853** | 10.41 | 61 | 673566** | 10.14 | 51 | 991896** | 10.23 | 71 | 1016062** | 10.09 |
| IPCA5 | 39 | 554526** | 7.70 | 59 | 531312** | 7.74 | 49 | 764772** | 7.58 | 69 | 784367** | 7.57 |
| IPCA6 | 37 | 434503** | 5.72 | 57 | 366817** | 5.16 | 47 | 626832** | 5.96 | 67 | 590757** | 5.54 |
| IPCA7 | 35 | 304589** | 3.79 | 55 | 261755** | 3.55 | 45 | 435330** | 3.96 | 65 | 393624** | 3.58 |
| IPCA8 | 33 | 262829** | 3.09 | 53 | 200749** | 2.63 | 43 | 350276** | 3.05 | 63 | 291948** | 2.57 |
| IPCA9 | 31 | 196967** | 2.17 | 51 | 150720** | 1.9 | 41 | 257790** | 2.14 | 61 | 217117** | 1.85 |
| IPCA10 |  |  |  |  |  |  | 39 | 114102** | 0.90 | 59 | 118716** | 0.98 |
| error | 720 | 60853 | 3.49 | 1080 | 55228 | 3.00 | 1520 | 49684 | 3.39 | 2280 | 45089 | 2.91 |

### 3.2 Higher-Order Additive Main effect and Multiplicative Interaction (HO-AMMI)

Table 2 presents the HO-AMMI ANOVA for simulated peanut pod yield data set from four different experiments. Unlike AMMI model which computes only additive effects of genotype and environment and multiplicative interaction effect of GEI; HO-AMMI model computes direct effects of genotype (G), location (L) and year (Y) and multiplicative effect of all possible interactions effect such as genotype*location interaction (GLI), genotype*year interaction (GYI), location*year interaction (LYI) and genotype*location*year interaction (GLYI). Influence of G, L and Y was significant on pod yield of peanut among all four experiments. Significant influence of interaction effects such as LYI, GLI, GYI and GLYI were observed for pod yield among all four experimental data sets. Location explained maximum variation in pod yield which ranged from 54.6% (experiment 3) to 62.2% (experiment 2) followed by location*year interaction (LYI), genotype*location interaction (GLI), genotype*location*year interaction (GLYI), genotype (G) and genotype*year interaction (GYI) among all four experiments.

**Table 2:**
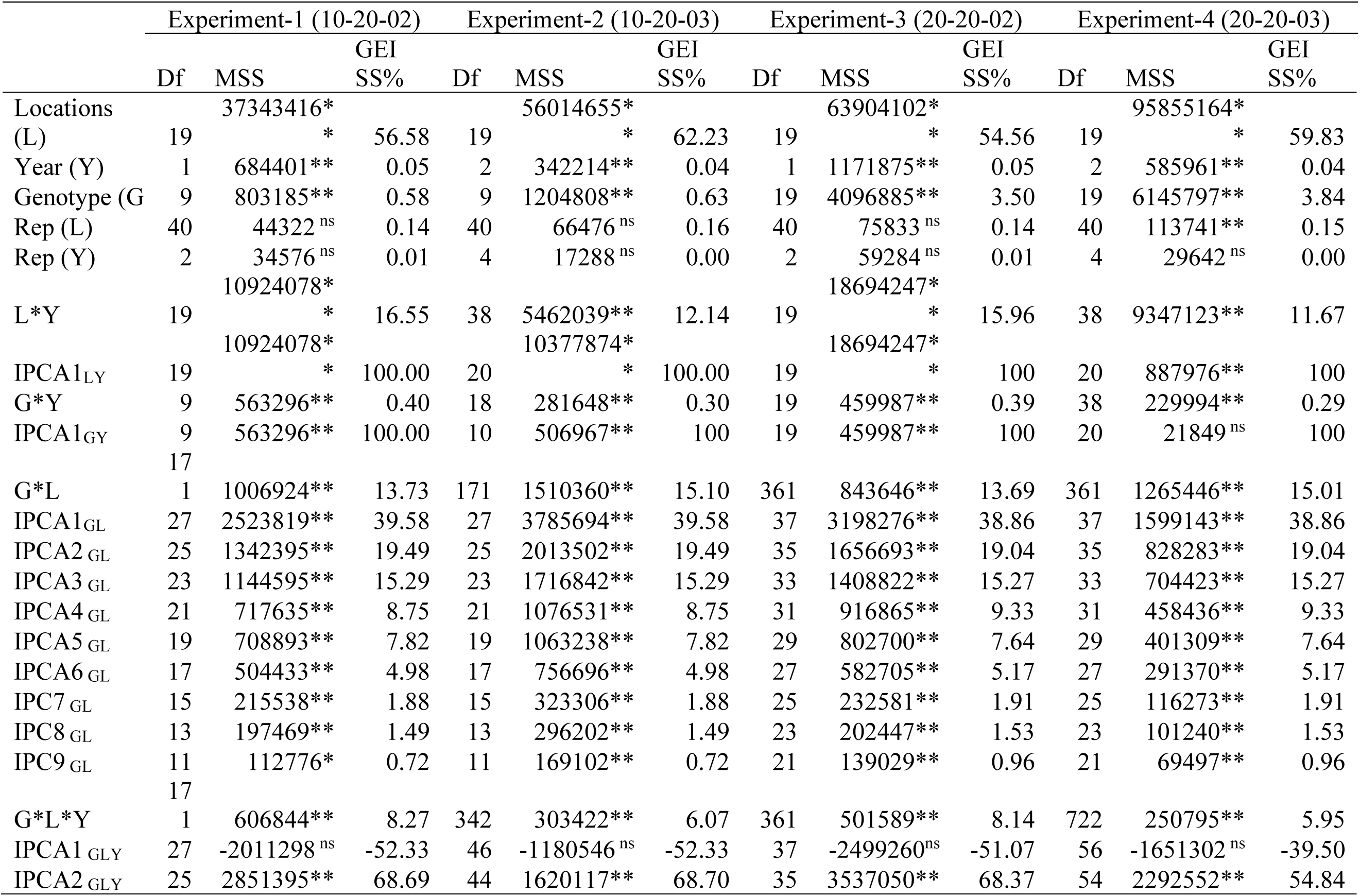
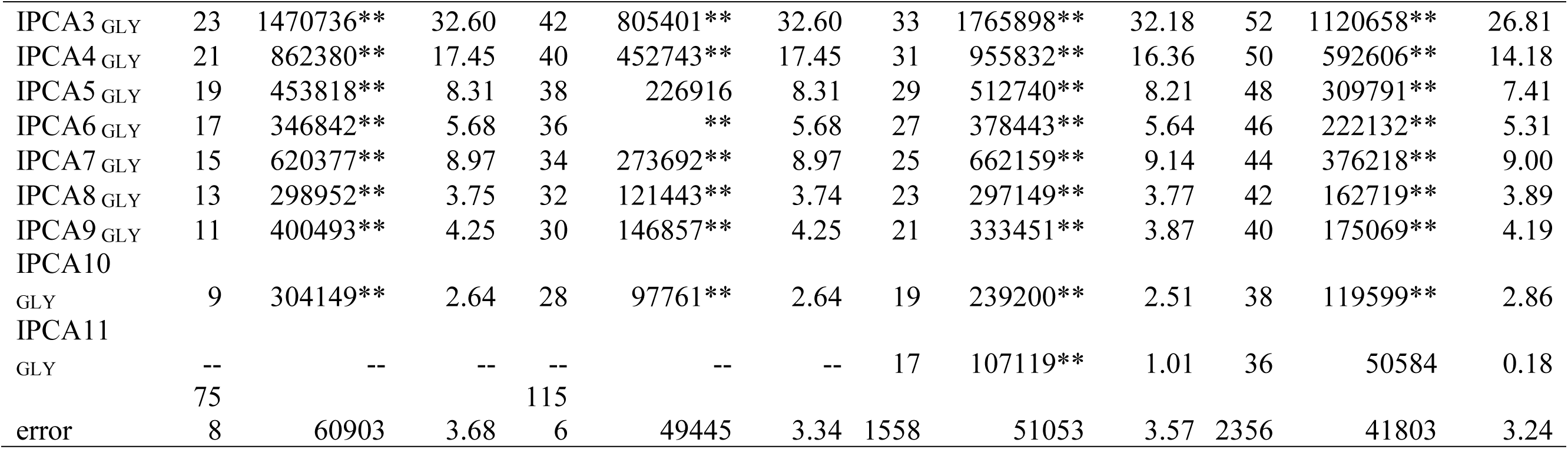
Higher order AMMI (HO-AMMI) analysis of variance for studying genotype-location-year-interactions (GLYI) under different combinations of genotype, location and year

LYI had one interaction principal components (IPCA1_LY_) which was significant among all four experiments whereas GYI had one interaction principal components (IPCA1_GY_) which was significant in experiments 1 to 3 and non-significant in experiment 4. GLI had nine significant IPCAs among all four experiments explaining 100% of GLI sum of squares. Percent contribution of IPCA1-9_GY_ towards GLI was same between experiments 1 and 2 and experiments 3 and 4 even though genotypic mean in experiment-1 with two year trial was different from experiment-2 with three year trial (Table S2 and S3), similarly genotypic mean of 20 genotypes in experiment-3 with two year trial was different from experiment-4 with three year trial (Table S4 and S5). This indicates that GLI will provide an accurate information about performance of genotypes over wide variety of locations irrespective of number of years of evaluation. GLYI had ten significant IPCAs in experiments 1 and 2 and eleven significant IPCAs in experiments 3 and 4 which all together explained 100% variation in GLYI sum of squares. In GLI or GYI or LYI wherein interaction components are arranged in the order of variation explained with first component explaining maximum variation and succeeding components explaining low variation compared to previous ones and these interaction components follow χ^2^ distribution (Gollob 1968). But in case of GLYI, interaction components follow noncentral χ^2^ distribution (Gollob 1968) and as a result component are not arranged in the descending order. In the present study variation explained by IPCA1_GLY_ was very low with negative sum of squares and non-significant, IPCA2_GLY_ explained highest variations followed by IPCA3_GLY_, IPCA4_GLY_, IPCA5_GLY_ and others in descending order. As IPCA1_GLY_ was negative and non-significant they need not be considered as a part of common variance (Lorenzo-Seva 2013).

### 3.3 Comparison of HO-AMMI vs AMMI

Major difference between HO-AMMI and AMMI is the estimation of interactions which are confounded within GEI. In experiment-1, GEI (22.4%) of AMMI was partitioned by HO-AMMI into GLI (13.73%), GYI (0.4%) and GLYI (8.27%). In experiment-2, GEI (21.5%) was partitioned into GLI (15.1%), GYI (0.3%) and GLYI (6.07%). Similarly, in experiment-3 and 4, GEI (22.1 and 21.24%) was partitioned into GLI (13.69 and 15.1%), GYI (0.39 and 0.29%) and GLYI (8.14 and 5.95%). This indicates that HO-AMMI was able to remove the confounding effect of GYI and GLYI on GLI. Earlier reports have also indicated that genotypic (V_G_) and genotype*location (V_GL_) variances, were confounded by genotype*year (V_GY_) and genotype*year*location (V_GYL_) variances, respectively (Holland and Nyquist 2010; Arief et al. 2015). Hence, HO-AMMI model presented in this study provides an accurate estimation of genotype*location (V_GL_) variances without the confounding effect of genotype*year (V_GY_) and genotype*year*location (V_GYL_) variances.

Both AMMI and HO-AMMI models behave differently when genotypes or years of evaluation were increased. In AMMI model, with increase in number of years of evaluation variation explained by G increased whereas that of GEI reduced. In HO-AMMI model, increase in number of years of evaluation increased percent contribution of GLI and reduced percent contribution of LYI and GLYI towards total variation. Increase in number of genotypes influenced mainly additive effects whereas interaction effects remained unchanged in both AMMI and HO-AMMI models. In AMMI model, genotype main effect increased and environment main effect decreased and GEI remained unchanged. In HO-AMM model, genotype main effect increased, location main effect decreased whereas year main effect and all other interaction effects remained unchanged. Differences in interaction effect of GEI and GLI over the years also indicated that in MET trials genotypes should be evaluated in as many number of years as possible in order to improve the accuracy of interaction effects in both AMMI and HO-AMMI models. Arief *et al*. (2019) have also indicated that the accuracy of V_GY_ and V_GYL_ estimates are affected by the number of years of evaluation. Another noticeable difference between AMMI and HO-AMMI models is difference in the total amount of variation explained by main effects and interaction effects. In HO-AMMI model percent variation explained by multiplicative components is higher in comparison to AMMI model.

### 3.4 Comparison of genotype ranking

Table S6 presents ranking of genotypes based on SSI for interactions such as GLI, GYI, GLYI and GLI+GYI+GLYI of HO-AMMI; GEI of AMMI model and field ranking. There were clear differences in genotype ranking between different interactions of HO-AMMI, AMMI and field ranking among all four experimental data sets. Experiments 1 and 2 have similar sets of genotypes and locations but differ in the number of years of evaluation and similarly with respect to experiments 3 and 4. Ranking of genotypes on the basis of GLI and GYI was same between experiments 1 and 2 and between experiments 3 and 4 whereas ranking of genotypes based on GLI+GYI+GLYI, field rank and AMMI differed between different years of testing. This indicates that when AMMI is applied ranking of genotypes vary with different years of evaluation which makes selection of genotypes cumbersome. But GLI interaction evaluated by HO-AMMI provides accurate ranking of genotypes irrespective of number of years of evaluation as confounding effects of GYI and GLYI have been removed. This further corroborates the observations made by earlier reports that V_G_ and V_GL_ were confounded by V_GY_ and V_GYL_ (Holland and Nyquist 2010; Arief et al. 2015).

### 3.5 Association among different ranking

Correlation analysis was performed between field ranking and interaction components of AMMI (GEI) and HO-AMMI models (GLI, GYI, GLYI, GLI+GYI+GLYI) and correlation values are presented in Table 3. In experiment 1, GLI and GLI+GYI+GLYI of HO-AMMI and AMMI had significant positive association with field ranking. In experiment 2, GLI, GYI and AMMI had significant positive association with field ranking and GLI had significant positive association with GLI+GYI+GLYI and AMMI ranking. In experiment 3 and 4, field ranking had significant association with GLI, GYI, GLYI, GLI+GYI+GLYI and AMMI; GLI had significant association with GLYI, GLI+GYI+GLYI and AMMI. In all four experiments, it was observed that field rank had higher correlation values with GLI compared to GYI, GLYI, GLI+GYI+GLYI and AMMI. Linear regression analysis was performed between field ranking with GL and AMMI and their scatter plots with R^2^ values are presented in Fig 1. R^2^ values of GLI with field rank was higher than AMMI with field rank. Root mean square error (RMSE) values were calculated for GLI of HO-AMMI and GEI of AMMI models for all four experiments and are presented in Fig 2. RMSE values were low for GLI among all four experiments. High correlation, high R^2^ values and low RMSE values for GLI of HO-AMMI when compared to GEI of AMMI indicates that GLI from HO-AMMI model accurately predicts genotype ranking compared to AMMI model.

**Table 3:**
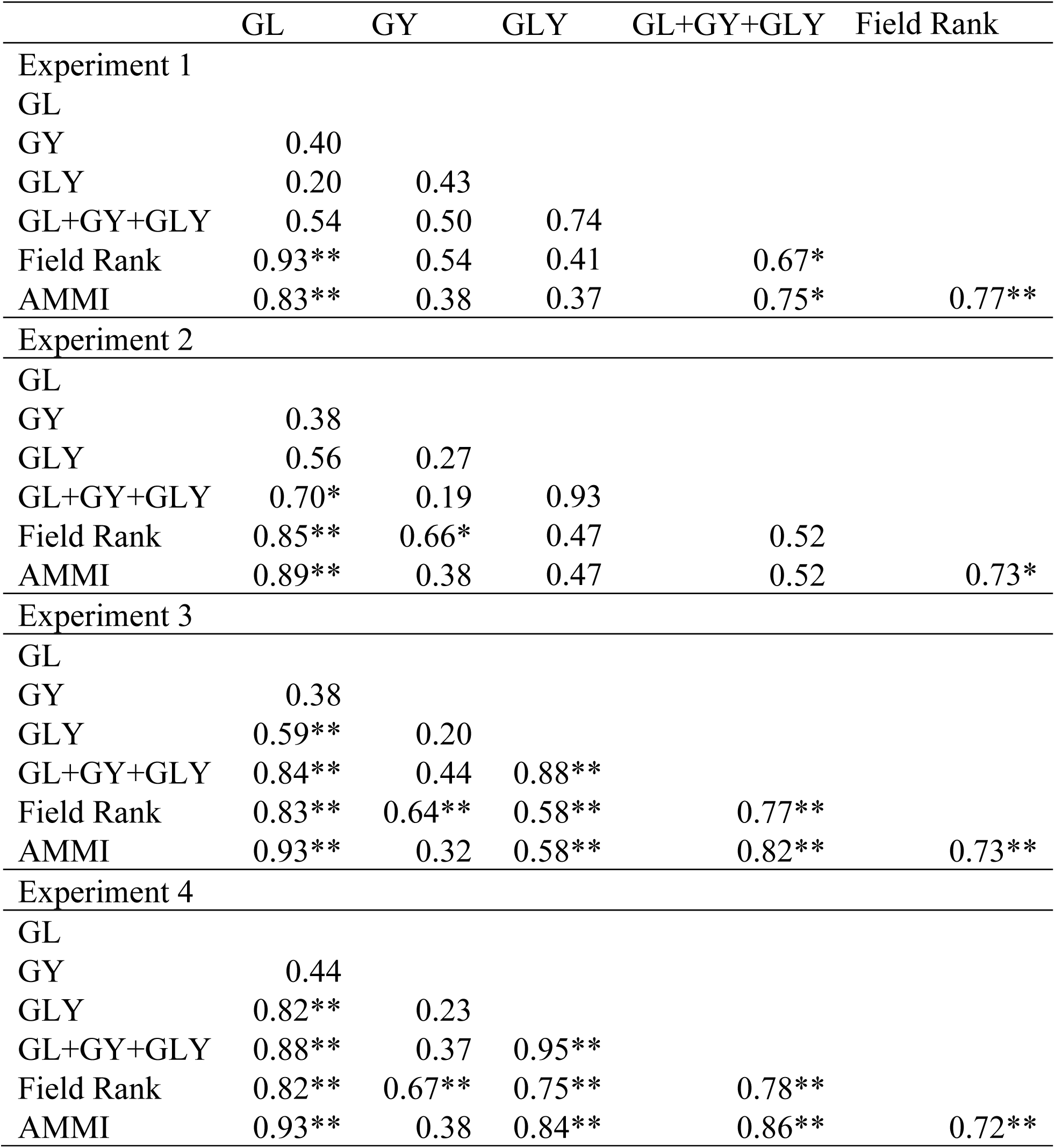
Linear regression analysis (R^2^) performed between GL, GY, GLY, GL+GY+GLY interactions of HO-AMMI and GEI interaction of AMMI with field ranking.

|  | GL | GY | GLY | GL+GY+GLY | Field Rank |
| --- | --- | --- | --- | --- | --- |
| Experiment 1 |  |  |  |  |  |
| GL |  |  |  |  |  |
| GY | 0.40 |  |  |  |  |
| GLY | 0.20 | 0.43 |  |  |  |
| GL+GY+GLY | 0.54 | 0.50 | 0.74 |  |  |
| Field Rank | 0.93** | 0.54 | 0.41 | 0.67* |  |
| AMMI | 0.83** | 0.38 | 0.37 | 0.75* | 0.77** |
| Experiment 2 |  |  |  |  |  |
| GL |  |  |  |  |  |
| GY | 0.38 |  |  |  |  |
| GLY | 0.56 | 0.27 |  |  |  |
| GL+GY+GLY | 0.70* | 0.19 | 0.93 |  |  |
| Field Rank | 0.85** | 0.66* | 0.47 | 0.52 |  |
| AMMI | 0.89** | 0.38 | 0.47 | 0.52 | 0.73* |
| Experiment 3 |  |  |  |  |  |
| GL |  |  |  |  |  |
| GY | 0.38 |  |  |  |  |
| GLY | 0.59** | 0.20 |  |  |  |
| GL+GY+GLY | 0.84** | 0.44 | 0.88** |  |  |
| Field Rank | 0.83** | 0.64** | 0.58** | 0.77** |  |
| AMMI | 0.93** | 0.32 | 0.58** | 0.82** | 0.73** |
| Experiment 4 |  |  |  |  |  |
| GL |  |  |  |  |  |
| GY | 0.44 |  |  |  |  |
| GLY | 0.82** | 0.23 |  |  |  |
| GL+GY+GLY | 0.88** | 0.37 | 0.95** |  |  |
| Field Rank | 0.82** | 0.67** | 0.75** | 0.78** |  |
| AMMI | 0.93** | 0.38 | 0.84** | 0.86** | 0.72** |

**Fig 1:**
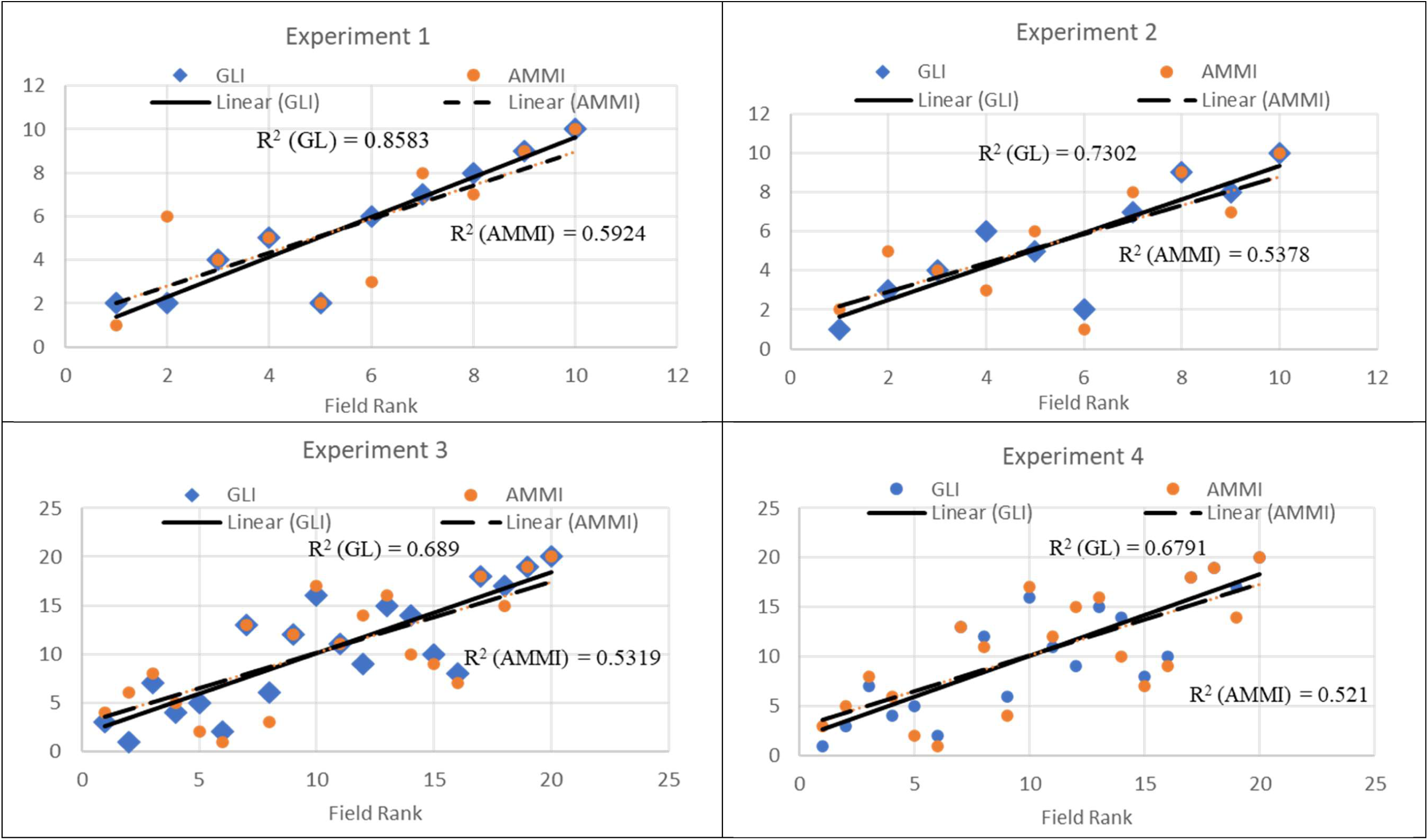
Linear regression of genotype*location interaction (GLI) from HO-AMMI model and AMMI model with field rank under different experimental conditions.

**Fig 2:**
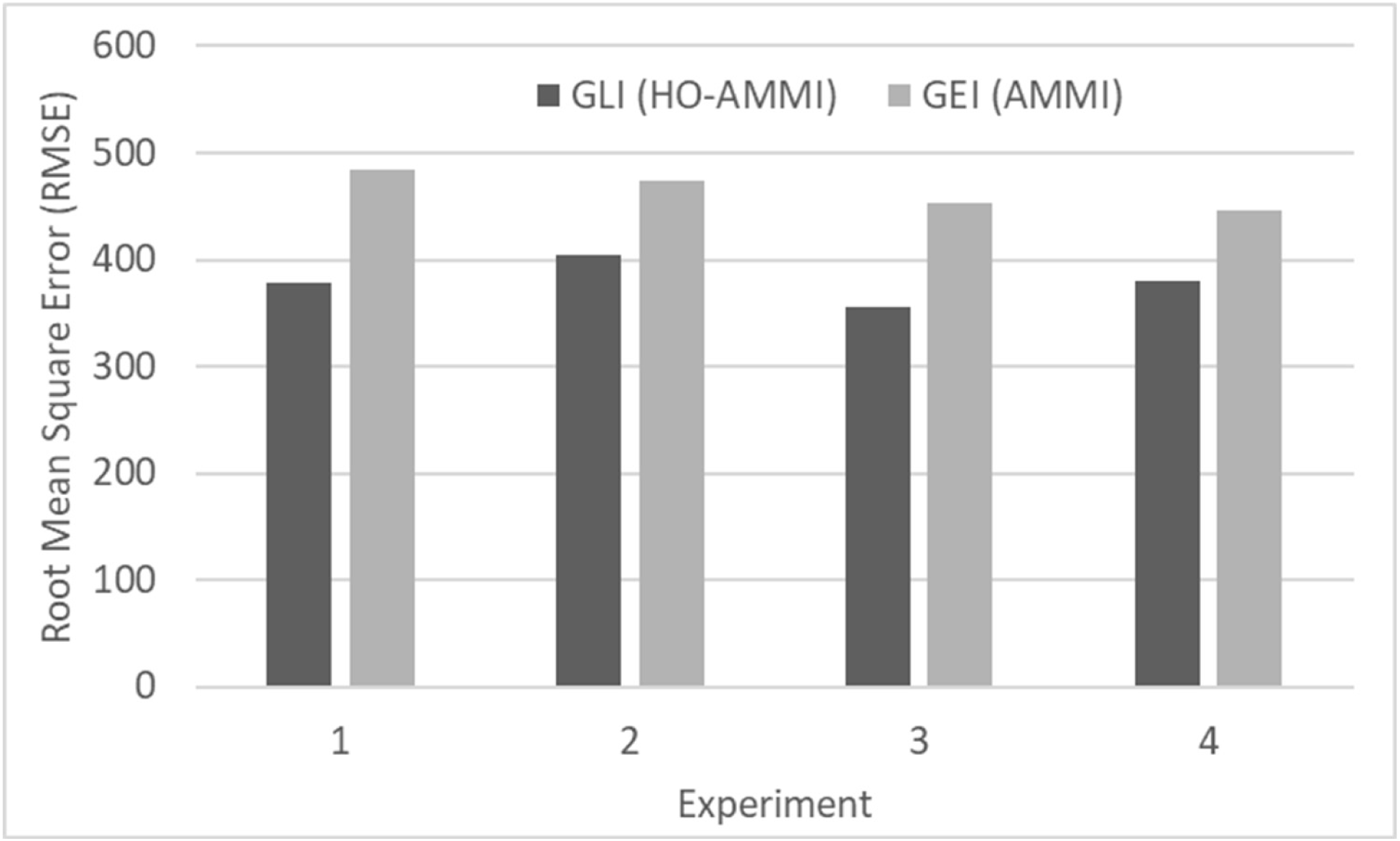
Root Mean Square Error (RMSE) for GLI from HO-AMMI and GEI from AMMI model under different experiments

### 3.6 Biplots comparison of HO-AMMI and AMMI models

The IPCA1 versus IPCA2 biplot of AMMI model explain the magnitude of interaction of each genotype and environment. The genotypes and environments that are farthest from the origin being more responsive. Genotypes and environments that fall into the same sector interact positively; negatively if they fall into opposite sectors (Osiru et al. 2009). A genotype showing high positive interaction in an environment obviously has the ability to exploit the agro-ecological or agro-management conditions of environment and is therefore best suited to that environment. Purchase (1997) pointed out that the closer the genotypes score to the center of the biplot, the more stable they are. But when environment consists of both locations and years selection of genotypes for particular location becomes difficult in AMMI model as GLI effect is masked by GYI and GLYI interaction effects. Confounding effect of year is overcome in HO-AMMI model wherein GEI is partitioned into GLI, GYI and GLYI. Biplots for HO-AMMI were generated based on GLI as they had high R^2^ values with field ranking compared to other models. Then biplots of GLI from HO-AMMI model was compared with AMMI model.

Fig 3a and 3b represents GLI based HO-AMMI model for experiment 1 and 2 respectively. Distribution of genotype points in both the experiments revealed that the genotype-1 and 10 are close to the origin with minimal GL effect whereas remaining eight genotypes are scattered away from the origin and are affected by GLI effect. The genotype-2 had positive interaction with location E13, hence exhibited specific adaptation to that environment. Genotype ‘10’ displayed positive interaction with locations E8, E12, E19 and E20; genotype-3 with location E10, E12 and E8; genotype-6 with location E4; genotype-1 with locations E1, E9 and E18; genotypes-4 and ‘5’ with locations E6, E1, E7 and E11 and genotype-9 with locations E3, E14, E15 and E16.

**Fig 3:**
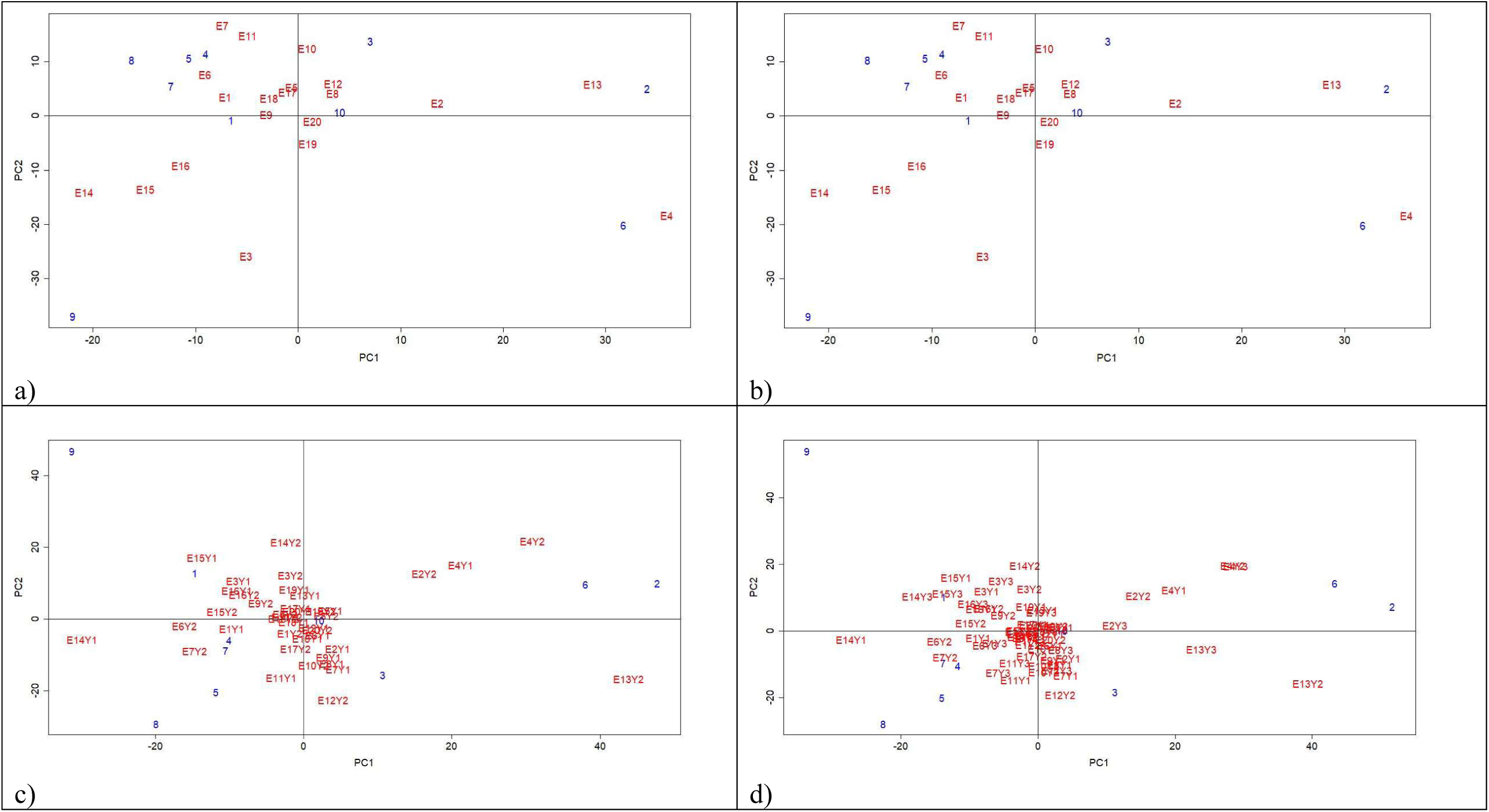
Biplot showing the effect of primary and secondary interaction principal components (PC1 and PC2) respectively for genotype*location interaction (GLI) for HO-AMMI and genotype*environment interaction for AMMI model under experiment 1 and 2: a) HO-AMMI experiment 1, b) HO-AMMI experiment 2, c) AMMI experiment 1 and d) AMMI experiment 2

Fig 3c and 3d represents AMMI model for experiment 1 and 2 respectively. Genotypes-10 is close to the origin with minimal GEI effect in both the experiments and remaining genotypes were sensitive to environment. In experiment-1 genotypes-4, 5 and 7 had had specific adaptation with environments E6Y2, E1Y1 and E7Y2; genotype-3 with E7Y1, E9Y1, E8Y1 and E2Y1; genotype-1 with E15Y1 and E3Y1. In experiment-2 genotypes-4 and 7 were adapted to environments E6Y2, E6Y3, E1Y1 and E7Y2; genotype-3 adapted to E7Y1, E12Y2 and E2Y1; genotype-1 with E14Y3, E15Y3 and E15Y1. Genotype-10 with maximum number of genotypes whereas genotypes-2, 8 and 9 suitable for none of the environments in both the experiments.

Fig 4a and 4b represents IPCA1vs IPCA2 biplot of GLI based HO-AMMI model for experiment 3 and 4 respectively. Distribution of genotype points in the biplot of both the experiments revealed that the genotype-1, 10, 11 and 20 are close to the origin, with minimal GLI effect whereas remaining sixteen genotypes are scattered away from the origin and are affected by GLI effects. The genotype-2 had positive interaction with location E13, hence exhibited specific adaptation to that environment. Similarly genotype-12 displayed positive interaction with locations E2; genotype-3, 10, 13 and 20 with location E20, E12 and E8; genotype-6 with location E4; genotype-1 and 11 with locations E5, E9, E17 and E18; genotypes-4, 5, 7, 8, 17, 18 with locations E6, E9, E7, E8, E14, E15, E17 and E18 and genotype-9 and 19 with locations E3, E14, E15 and E16.

**Fig 4:**
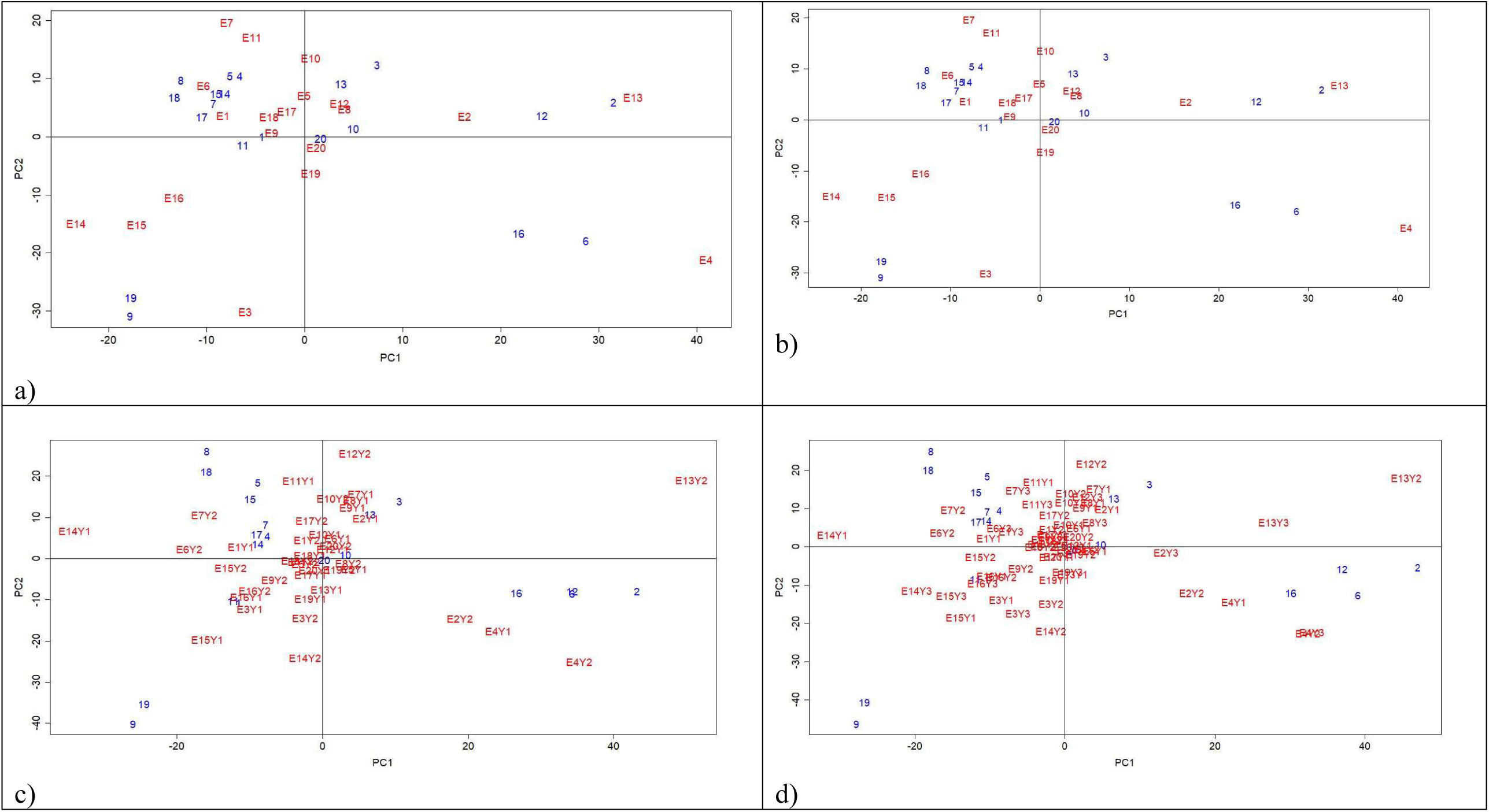
Biplot showing the effect of primary and secondary interaction principal components (PC1 and PC2) respectively for genotype*location interaction (GLI) for HO-AMMI and genotype*environment interaction for AMMI model under experiment 3 and 4: a) HO-AMMI experiment 3, b) HO-AMMI experiment 4, c) AMMI experiment 3 and d) AMMI experiment 4

Fig 4c and 4d represents IPCA1vs IPCA2 biplot of AMMI model for experiment 3 and 4 respectively. Genotypes-10 and 20 are close to the origin with minimal GEI effect in both the experiments and remaining genotypes were scattered around the biplot and are sensitive to environmental effects. In experiment-3 genotypes-4, 5, 7, 14, 15 and 17 had had specific adaptation with environments E6Y2, E1Y1, E7Y2 and others; genotype-3 ad 13 with E7Y1, E9Y1, E8Y1, E2Y1 and others; genotype-1 and 11 with E15Y1, E3Y1, E16Y2 and others. In experiment-4 genotypes-4, 5, 7, 14, 15 and 17 were adapted to environments E6Y2, E6Y3, E1Y1, E7Y2 and others; genotype-3 and 13 adapted to E7Y1, E12Y2, E2Y1, E8Y3 and others; genotype-1 and 11 with E14Y3, E15Y3, E15Y1 and others. Genotype-10 and 20 with maximum number of genotypes. Genotypes 8, 9, 18, 19 were suitable for none of the environments in both the experiments.

Association and distribution of genotypes with locations in HO-AMMI model was similar between experiments 1 and 2 and between 3 and 4 irrespective of number of years of evaluation. Whereas in AMMI model though genotypic distribution in biplot was almost same between experiments (experiments 1 vs 2 and experiment 3 vs 4) distribution of environments varied. In AMMI model total number of environments is a product of number of locations and number of years of evaluation. So, a trial with 20 locations and 2 years will have 40 environments and number of environments increases with the increase in locations and years. Whereas in HO-AMMI model irrespective of number of years of evaluation GLI biplot will produce similar results. Also, selection of genotypes for target location is easy in HO-AMI model as GLI biplot is based on genotypes and locations only without the year effect. Whereas in AMMI model year effect is masking the location effect making genotypic selection difficult.

## 4. Conclusion

HO-AMMI model was able to partition the GEI into several two-way and three-way interactions and accurately calculate GLI without the confounding effect of GYI and GLYI. HO-AMMI model was able to provide accurate ranking the genotypes for a location irrespective of number of years of evaluation. Also, GLI biplots from HO-AMMI model depicted very clear picture about the association of genotype with target location which was missing in AMMI models. Hence, HO-AMMI model would help the breeder to identify stable high yielding genotype for a target location without the confounding effect of years.

## Declaration

The authors declare no conflict of interest.

## Abbreviations

AMMI: Additive main effects and multiplicative interaction
HO-AMMI: Higher-order Additive main effects and multiplicative interaction
GEI: Genotype*environment interactions
GLI: Genotype*location interaction
GYI: Genotype*year interaction
GLYI: Genotype*location*year interaction
LYI: Location*year interaction
MET: Multi-environment trials

## Acknowledgements

Authors would like to thank ICAR and Director ICAR-DGR for supporting this project.

## Author Contributions

A.B.C. and B.S.K conceptualized the experiment, P.K, K.R, N.K and G.K generated the hypothetical data; A.B.C and A.F.R analysed the results; A.B.C and R.T wrote the manuscript; and All authors reviewed the manuscript.

